# Chromosomal DNA sequences of the Pacific saury genome: versatile resources for fishery science and comparative biology

**DOI:** 10.1101/2023.10.16.562003

**Authors:** Mana Sato, Kazuya Fukuda, Mitsutaka Kadota, Hatsune Makino-Itou, Kaori Tatsumi, Shinya Yamauchi, Shigehiro Kuraku

**Affiliations:** Molecular Life History Laboratory, National Institute of Genetics, Mishima, Japan; Laboratory of Reproductive Physiology of Aquatic Organisms, School of Marine Biosciences, Kitasato University; Laboratory for Phyloinformatics, RIKEN Center for Biosystems Dynamics Research, Kobe, Japan; Fukushima Marine Science Museum (Aquamarine Fukushima), 50 Tatsumi-cho, Onahama, Iwaki, Fukushima, 971-8101, Japan; Department of Genetics, Sokendai (Graduate University for Advanced Studies), Mishima, Japan

**Author notes:** These authors contributed equally (Mana Sato, Kazuya Fukuda, Mitsutaka Kadota, Hatsune Makino-Itou, Kaori Tatsumi). Correspondence. Yata 1111, Mishima, Shizuoka 411-8540, Japan.

**Keywords:** Pacific saury, Beloniformes, chromosome-level genome assembly, aquaporin, Hi-C, genome size

## Abstract

Pacific saury (*Cololabis saira*) is a commercially important small pelagic fish species in Asian. In this study, we conducted the first-ever whole genome sequencing of this species, with single molecule, real-time (SMRT) sequencing technology. The obtained high-fidelity (HiFi) long-read sequence data, which amount to approximately 30 folds of its haploid genome size that was measured with quantitative PCR (1.17 Gb), were assembled into contigs. Scaffolding with Hi-C reads yielded a whole genome assembly containing 24 chromosome-scale sequences, with a scaffold N50 length of 47.7 Mb. Screening of repetitive elements including telomeric repeats was performed to characterize possible factors that need to be resolved towards ‘telomere-to-telomere’ sequencing. The larger genome size than in medaka, a close relative in Beloniformes, is at least partly explained by larger repetitive element quantity, which is reflected in more abundant tRNAs, in the Pacific saury genome. Protein-coding regions was predicted using transcriptome data, which resulted in 22,274 components. Retrieval of Pacific saury homologs of aquaporin (AQP) genes known from other teleost fishes validated high completeness and continuity of the genome assembly. These resources are available at https://treethinkers.nig.ac.jp/saira/ and will assist various molecular-level studies in fishery science and comparative biology.

## 1. Introduction

The Pacific saury or mackerel pike (Fig. 1; *Cololabis saira* Brevoort, 1856) is a carnivorous, marine thermophilic shallow-water dweller and is one of the most popular dominant epipelagic nekton species in the North Pacific. Inside the taxon Beloniformes, the lineages of the Pacific saury and Japanese medaka (*Oryzias latipes*) diverged from each other approximately 74 million years^1^. Analyses based on the comparative molecular biology and genomics are expected to address anatomical and behavioral questions. However, the lack of molecular sequence information is an obstacle.

**Figure 1.**
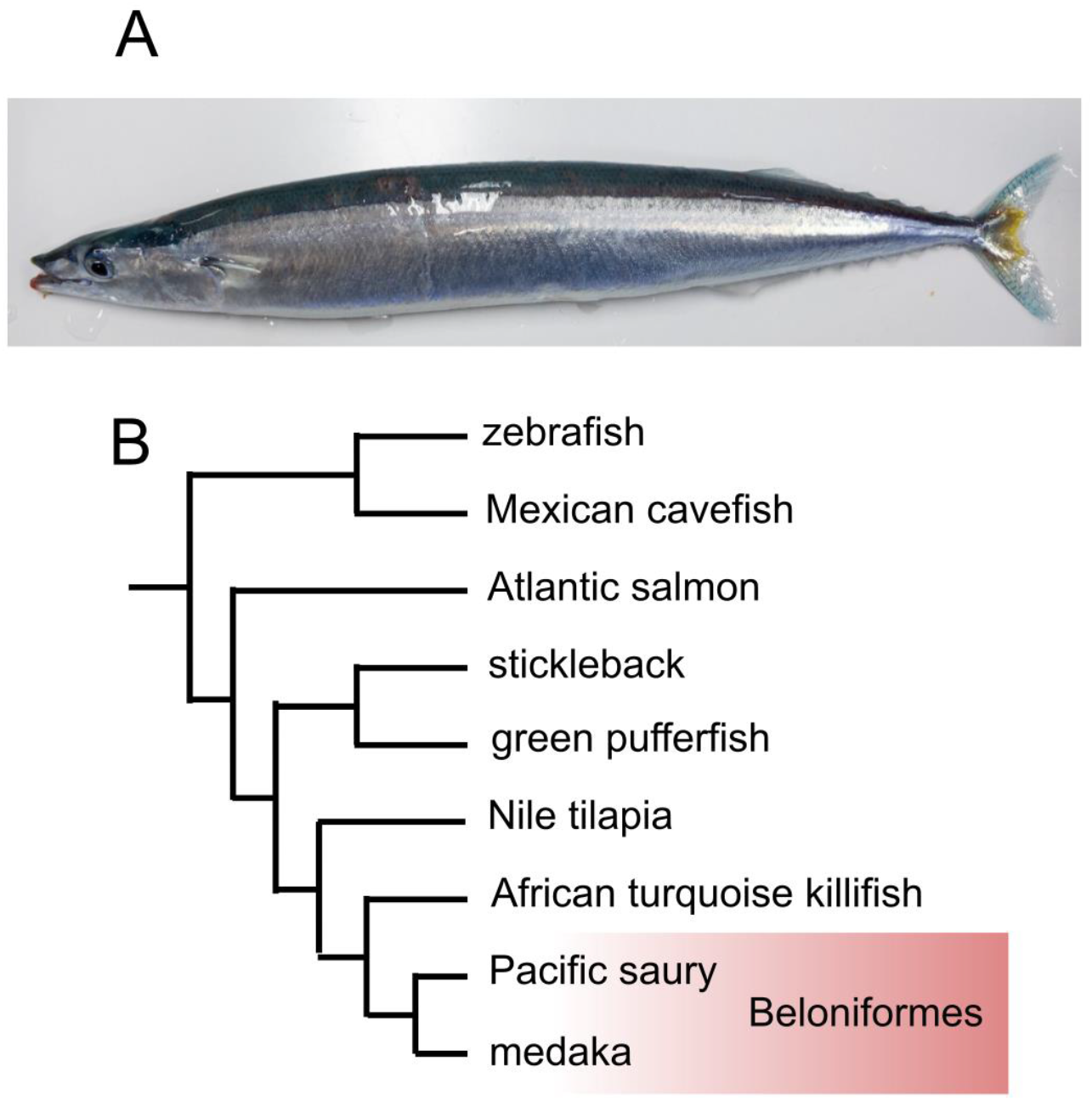
Pacific saury. A) Adut fish of approximately 29.5 cm in fork length. B) Position of the Pacific saury in the phylogeny of selected teleost fish species. The phylogenetic position is based on molecular phylogeny presented in existing literature^2,3^.

For its importance in fisheries, molecular-level studies of the Pacific saury are severely lacking. As of May 2023, the number of entries of nucleotide and protein sequences in NCBI^4^ are 150 and 99, respectively, most of which are fragments of mitochondrial DNA and peptide sequences encoded by them. In the Beloniformes that the Pacific saury belongs to, 337 species has been described as of May, 2023 according to the Eschmeyer’s Catalog of Fishes^5^, among which the whole-genome sequences of the nuclear DNA has been sequences and made publicly available only for nine species including as many as seven species that belongs to medaka fish genus *Oryzias*.

The nuclear DNA content (or genome size) of the Pacific saury has not been reported to date, while its karyotype was reported to be 2n = 42^6^. Importantly, this karyotype report lacked a detailed documentation of the karyotypic variation based on the observation of metaphase spreads of chromosomes prepared from multiple cells. Furthermore, the reported diploid number of chromosomes, namely 42, is distinct from that of most of the close relatives of this group, namely 48 or 50^7^, which requires reevaluation.

To build a basis of molecular-level research, in this study, we demonstrated a modern art of whole genome sequencing employing the high-fidelity long-read technology and obtained a chromosome-scale genome assembly by means of Hi-C scaffolding. Our product provides a fundamental resource for evidence-based analysis of fishery science and comparative biology.

## 2. Materials and Methods

### 2.1. Animals

The male individual of the Pacific saury used in this study (ToLID; fColSai1) was sampled in May 2022 from the exhibition tank of the public aquarium Aquamarine Fukushima, in Iwaki, Japan. Animal handling and sample collections at the aquaria were conducted by veterinary staff without restraining the individuals, in accordance with the Husbandry Guidelines approved by the Ethics and Welfare Committee of Japanese Association of Zoos and Aquariums.

### 2.2. Gonadal histology

Histological analysis was performed using the gonad of the individual sampled above in May 2022 for genome sequencing. Also, tissues of a male and a female collected in June 2022 were fixed in Bouin’s fixative and used for histological observations as a reference to the gonadal structure of each sex. All tissues were embedded in paraffin (Leica Biosystems, Nussloch, Germany), sectioned in 5 μm, and stained with Mayer’s hematoxylin (Fujifilm Wako Pure Chemical Co., Osaka, Japan) and eosin Y (Fujifilm Wako Pure Chemical Co.).

### 2.3. Genome sequencing and assembly

High molecular weight DNA was extracted from blood cells using a NucleoBond AXG column (Macherey-Nagel, Düren, Germany), which was followed by purification with phenol-chloroform. The concentration of the extracted DNA was measured with Qubit 4 (ThermoFisher, MA, USA), and their size distribution was first analyzed with TapeStation 2100 (Agilent Technologies, CA USA) to ensure high integrity and later analyzed with pulse-field gel electrophoresis on CHEF DR-II (BioRad, CA, USA) to ensure the size range between 20 kb to 100 kb. The DNA was fragmented with g-TUBE (Covaris, MA, USA) and size-selected with BluePippin according to the official protocol. An SMRT sequence library was constructed with an SMRTbell Express Template Prep Kit 2.0 (Pacific Biosciences, CA, USA) and was sequenced in a single 8M SMRT cell on a PacBio Sequel IIe system (Pacific Biosciences). The sequencing output was processed to generate circular consensus sequences (CCS) to obtain a total of 33.7 Gb HiFi sequence reads. From these reads, adapter sequences were removed using the program HiFiAdapterFilt^8^. The obtained HiFi sequence reads were assembled using the program hifiasm v0.16.1^9^ with its default parameters. The obtained contigs were subjected to haplotig purging by using the program purge_haplotigs^10^ with the options ‘-l 5 -m 23 -h 45’.

### 2.4. Hi-C data production and genome scaffolding

The obtained contigs were scaffolded using Hi-C read pairs obtained as follows. The Hi-C library was prepared using the muscle tissue of the individual fColSai1 used for genome sequencing, according to the iconHi-C protocol employing restriction enzymes DpnII and HinfI^11^, and it was sequenced on a HiSeq X sequencing platform (Illumina Inc., CA, USA). The obtained Hi-C read pairs were processed with the program Trim Galore! v0.6.8 (https://www.bioinformatics.babraham.ac.uk/projects/trim_galore/) specifying the options ‘--phred33 --stringency 2 --quality 30 --length 25 --paired’ and aligned to the HiFi sequence contigs with the program Juicer v1.6^12^, and using its results, HiFi sequence contigs were scaffolded with 3d-dna^13^ (version 201008) specifying the options ‘-m haploid -i 5000 --editor-repeat-coverage 15 -r 2’ to be consistent with the chromatin contact profiles. The continuity and completeness of the resultant genome assembly, designated as fColSai1.1, as well as the predicted protein-coding gene set (see below), was assessed with the webserver gVolante v2.0.0^14^ in which the pipeline BUSCO v5.1.2 is implemented^15^, consistently using the ortholog set ‘actinopterygii_odb10’ supplied with BUSCO. Especially, the completeness of the nucleotide sequences of the genome assembly was assessed with compleasm^16^ (formerly called minibusco) that is thought to achieve higher accuracy due to the use of miniprot^17^. The genome assembly was deposited in NCBI under the accession ID PRJNA1027303.

### 2.5. Annotation of non-coding sequences

The obtained chromosome-scale and other genomic sequences were subjected to the *de novo* detection of repetitive elements with RepeatModeler v2.0.4 with the -LTRStruct option. The detected repeat sequences were input in RepeatMasker v4.1.4 for repeat-masking in the sensitive mode (with the option ‘-s’). Simple tandem repeat detection was further reinforced by the use of the program tantan^18^. Putative transfer RNAs (tRNAs) were detected by the program tRNAscan-SE version 2.0.12 and the detected regions excluding those labelled as ‘pseudogenized’ were counted. Detection of the rDNA-containing regions was performed with the program barrnap version 0.9 (https://github.com/tseemann/barrnap) with an aid of the partial sequence entry spanning the Pacific saury rDNA locus (OP151180.1) and the human 45S pre-ribosomal sequence (NR_145819.1). Detection of canonical telomeric repeats (TTAGGG)n was performed with tidk version 0.2.31 (https://github.com/tolkit/telomeric-identifier).

### 2.6. Genome size estimation

The genome size was quantified with the qPCR-based protocol sQuantGenome^19^ (https://github.com/Squalomix/c-value) with the same DNA sample used for whole genome sequencing. Three protein-coding genes, *ACAT1*, *DLD*, and *RFC3* from the single-copy orthologous gene set CVG^20^ (Core Vertebrate Genes), whose transcript sequences (Supplementary Table S1) were identified in the transcriptome sequence assembly (see below). The primers used for qPCR are included in Supplementary Table S2. Genome size in nucleotide base pairs was calculated based on the formula^21^ that the number of base pairs corresponds to mass in pg × 0.978 × 10^9^.

### 2.7. Transcriptome sequencing and assembly

For transcriptome data acquisition, an adult individual of unknown sex was sampled in July 2022 and was dissected. Total RNA extraction was performed for the heart, muscle, gill, liver, gut, and eye ball of this adult fish as well as a whole larvae body, using TRIzol reagent (ThermoFisher). After DNaseI digestion, strand-specific RNA-seq libraries were prepared using 150 ng of each of the extracted total RNAs (Supplementary Table S3), with Illumina Stranded mRNA Prep kit (Illumina, Cat. No. 20040534) and IDT for Illumina RNA UD Indexes Set A Ligation (Illumina, Cat. No. 20040553) according to its standard protocol unless stated otherwise below. Before the total volume PCR amplification was performed, we performed a preliminary PCR using a 1.5 μl aliquot of 10 μl DNA from the previous step, with KAPA Real-Time Library Amplification Kit (Kapa Biosystems, cat. No. KK2702). This demonstrated that the amplification of the products reached Standard 1 accompanying this kit between three and four PCR cycles, which instructed us to perform the full-volume PCR with three PCR cycles, introducing the minimal amplification (Supplementary Table S3). The libraries were sequenced in 0.6 lanes to obtain PE150 reads on a HiSeq X (Illumina). Quality control of the obtained fastq files for individual libraries was performed with FASTQC v0.11.5 (https://www.bioinformatics.babraham.ac.uk/projects/fastqc/).

The obtained sequence reads in the fastq files were processed with Trim Galore! v0.6.8 with the options --phred33--stringency 2--quality 30--length 25--paired’. The reads after adaptor trimming were assembled with the program Trinity v2.14.0 with the options --trimmomatic --SS_lib_type RF’. Among the resultant contig sequences, those matching PhiX, and mitochondrial DNA in the BLASTN v2.2.30+9 results executed with the options ‘-perc_identity 95’ were removed.

### 2.8. Molecular phylogenetics analysis

Protein sequences were collected from the NCBI and Ensembl databases, and their accession IDs used for the phylogenetic analysis are included in Supplementary Table S4. The deduced amino acid sequences were aligned with the MAFFT v7.505 using the L-INS-i method. The aligned sequences were trimmed with trimAl v1.4.rev15^22^ to remove unreliably aligned sites using the ‘-gappyout’ option. The maximum-likelihood tree was inferred with RAxML v8.2.12^23^ using the PROTCATWAG model, and for evaluating the confidence of the nodes, the rapid bootstrap resampling with 100 replicates was performed. Molecular phylogenetic tree employing the Bayesian framework was inferred with PhyloBayes v4.1c using the CAT-WAG-Γ model.

### 2.9. Gene prediction

Protein-coding genes were predicted with the program pipeline Braker3^24^ on the genome assembly sequences in which interspersed repeats and simple tandem repeats were soft-masked with RepeatModeler/RepeatMasker^25^ (see above) and tantan^18^. The gene prediction incorporated all peptide sequences registered in the file ‘odb11_vertebrata_fasta’ provided by the OrthoDB database^26^, as well as the output of paired-end RNA-seq read mapping with hisat2^27^.

## 3. Results and Discussion

### 3.1. Animal sampling and sex identification

An adult of the Pacific saury was sampled from the exhibition tank of Aquamarine Fukushima, and was subjected to histological analysis for identifying the sex and the stage of gonadal maturity, in parallel with high-molecular weight DNA extraction. Our histological observation of the Pacific saury individual revealed sexually mature testes, and it was judged to be a male (Fig. 2A-C). The testis showed a tubular-type structure (Fig. 2D) and had cysts containing various stages of germ cells; spermatozoa (Fig. 2E), spermatids (Fig. 2F), and spermatocytes (Fig. 2G). Abundant spermatozoa produced and stored in cysts close to the central cavity (Fig. 2E) indicates that this individual was sexually mature at the time of our sampling for genome sequencing.

**Figure 2.**
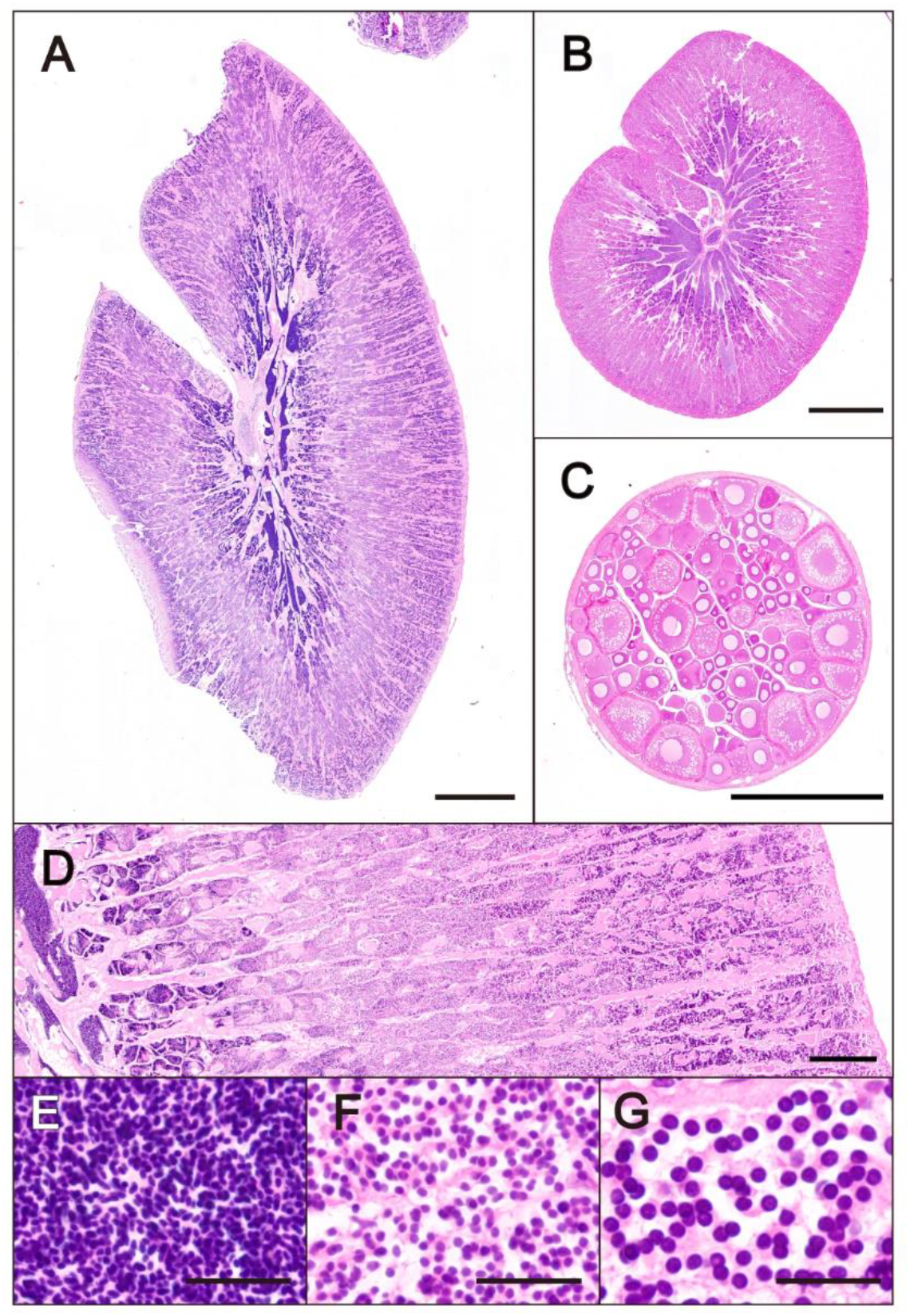
Histological sex identification. Gonadal cross sections of the individual whose genome was sequenced in this study (A) sampled in May 2022, and the reference male (B) and female (C) individuals sampled in June 2022, stained with hematoxylin and eosin staining (scale bar = 1 mm). (D-G) Detailed testis structure of the individual whose genome was sequenced in this study. Cysts are arranged depending on the developmental stages of the containing germ cells (D, scale bar = 200 μm). Germ cells located on the centrally, spermatozoa (E, scale bar = 20 μm); intermediate, spermatids (F, scale bar = 20 μm); outside, spermatocytes (G, scale bar = 20 μm).

### 3.2. Genome size estimation

The extracted genomic DNA was used for the estimation of nuclear DNA content with the method using quantitative PCR (qPCR)^19^. Three protein-coding genes included in Table 1 were selected from those contained in the transcript contigs (see Materials and Methods) and detected as single copy genes. We designed specific primers for these genes and prepared standard molecules for each of them by PCR. Using the standard molecules controlled with sequencing specificity and quantity, we performed qPCR and quantified the DNA amount that yields a haploid DNA molecule. This resulted in the haploid nuclear DNA content of 1.19 picogram (pg), which corresponds to 1.17 Gb.

**Table 1.** Summary of genome size estimation by qPCR.

| Target gene | DNA copy number<br>calculated by qPCR<sup>a</sup><br>(copies/μl DNA) | Estimated<br>C-value (pg) | Mean<br>C-value<br>(pg)<sup>b</sup> | Estimated haploid<br>genome size (Gbp)<sup>a</sup> |
| --- | --- | --- | --- | --- |
| ACAT1 | 5025.25 | 1.14 | 1.19 | 1.17 |
| DLD | 4335.21 | 1.32 |  |  |
| RFC3 | 5066.12 | 1.13 |  |  |

<sup>a</sup>A uniform amount of gDNA (5.71 ng/ μl) was used in qPCR.
<sup>b</sup>Mean value of the C-value estimated for the three individual genes.
<sup>c</sup>Genome size in nucleotide base pairs calculated according to the previously proposed formula (see Materials and Methods).

### 3.3. Genome sequencing and assembly

The extracted high molecular weight DNA was used for SMRTbell library and subjected to circular consensus sequencing (CCS) with single molecule, real-time (SMRT) sequencing technology (see Materials and Methods). This yielded 33.6 Gb high-fidelity long (HiFi) reads of approximately 30 folds of its haploid genome size (1.17 Gb). K-mer distribution in these reads suggested a heterozygosity of 1.87 % (Supplementary Fig. S1). These sequence reads were assembled into 1,555 contigs with a N50 length of 6.3 Mb. These contigs exhibited a percentage of duplicated one-to-one orthologs of 7.7 % and the total sequence length of 1.33 Gb that remarkably exceeded the genome size that was estimated independently (see above). We imputed these features to the relatively high heterozygosity (1.87%; see above) and purged redundant haplotigs. This resulted in 672 contig sequences with a reduced redundancy of one-to-one orthologs down to 1.9 % and a total sequence length of 1.15 Gb that is close to the genome size, 1.17Gb (Table 2).

**Table 2.** Genome assembly statistics in comparison with other teleost fishes.

| Species | *Cololabis saira* | *Oryzias latipes* | *Danio rerio* | *Gasterosteus aculeatus* | *Hirundichthys speculiger* | *Xenentodon cancila* |
| --- | --- | --- | --- | --- | --- | --- |
|  | Pacific saury | Medaka | zebrafish | three-spined stickleback | mirrorwing flying-fish | freshwater garfish |
| Assembly | fColSai1.1 | UT | GRCz11 | GAculeatus UGA version5 | GCA_018398805.1 | GCA_014839995.1 (fXenCan1.pri) |
| Total length (Mbp) | 1,146 | 790 | 1,679 | 472 | 1,043 | 736 |
| N50 scaffold length (Mbp) | 47.7 | 31 | 52 | 20 | 1 | 29 |
| Largest scaffold length (Mbp) | 57.6 | 38 | 78 | 34 | 9 | 36 |
| Number of scaffolds >10 Mbp | 24 | 24 | 25 | 22 | 0 | 24 |
| Number of scaffolds >1 Mbp | 24 | 25 | 25 | 22 | 259 | 25 |
| % of sum length of sequences >10 Mbp | 96.7 | 92.9 | 80.1 | 95.8 | 0.0 | 96.4 |
| % of sum length of sequences >1 Mbp | 96.8 | 93.1 | 80.1 | 95.8 | 8.5 | 97.1 |
| # Seqs | 758 | 920 | 1922 | 2937 | 3052 | 410 |
| # Gaps<sup>1</sup> | 896 | 491 | 1,8796 | 3,122 | 206 | 1,373 |
| % Gap<sup>1</sup> | 0.04 | 0.06 | 0.28 | 0.76 | 0.05 | 1.02 |
| % Complete ortholog<sup>2</sup> | 98.9 | 99.2 | 97.7 | 98.7 | 99.3 | 98.9 |
| % Complete+Fragment ortholog<sup>2</sup> | 99.3 | 99.5 | 99.1 | 99.4 | 99.7 | 99.3 |
| % Duplicate ortholog<sup>2</sup> | 1.84 | 0.4 | 17.0 | 1.6 | 3.9 | 0.2 |
| % Missing ortholog<sup>2</sup> | 0.66 | 0.5 | 0.9 | 0.6 | 0.3 | 0.6 |

<sup>1</sup>Gaps stand for a string of no shorter than five undetermined ('N') bases. <sup>2</sup>These scores stand for the percentage of reference orthologues identified in the input genome assembly and were obtained with the program compleasm (formerly, called minibusco) using the ortholog set Actinopterygii provided by the BUSCO developer (see Materials and Methods).

We prepared a Hi-C library using a combination of restriction enzymes DpnII and HinfI (see Materials and Methods) with the iconHi-C protocol (Kadota et al., 2020). This Hi-C library was sequenced on a short-read sequencing platform to obtain 142 million paired-end reads. Using these Hi-C reads, the contigs obtained above were scaffolded consistently with chromatin contact profiles (Fig. 3A). Subsequent manual curation by referring to the Hi-C contact map^28^ resulted in a whole genome assembly containing 24 chromosome-scale sequences, with a total nucleotide length of 1.146 Gb and a scaffold N50 length of 47.7 Mb. Of those chromosome-scale sequences, the longest sequence amounts to 57.6 Mb and the shortest one 27.1 Mb. Separately, we identified a 16,499 bp-long whole mitochondrial DNA sequence assembled from the obtained HiFi reads that was identical to the nucleotide sequence of the existing NCBI entry NC_003183.1 for the Pacific saury whole mitochondrial DNA. The whole genome assembly excluding this mitochondrial DNA sequence was designated fColSai1.1 and was used in downstream analyses. The genome assembly, as well as other sequence resources obtained in this study, are available as databases for BLAST^29^ searches at the original user-friendly webserver https://treethinkers.nig.ac.jp/saira/blast/ built with SequenceServer v2.0.0^30^.

**Figure 3.**
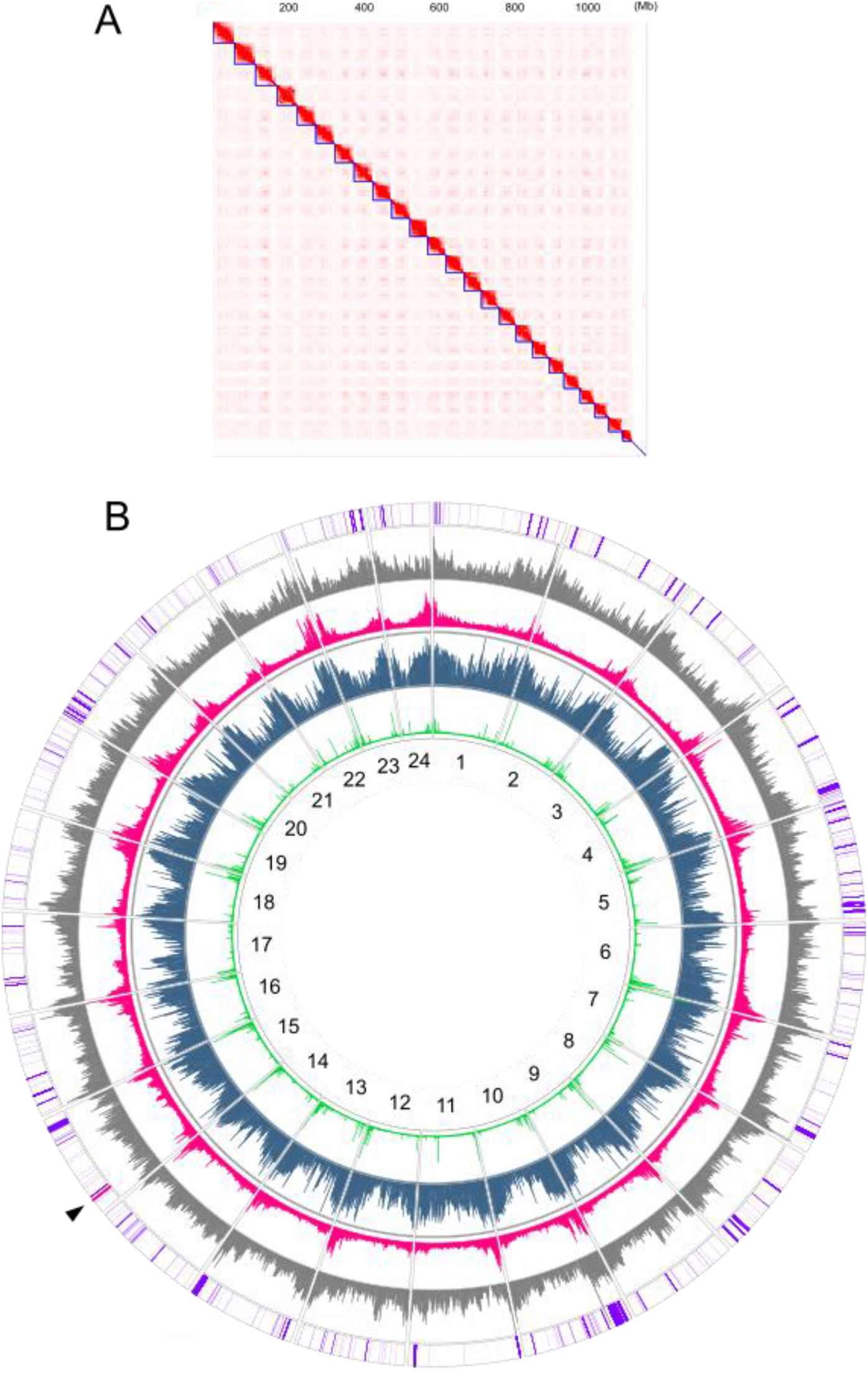
Pacific saury genome assembly. (A) Hi-C contact map. (B) Genomic landscape shown for individual chromosomes (black letters). This circular view shows the contents of GC nucleotides (grey; 40-50%), simple tandem repeats (red; 0-90%), interspersed repeats (blue; 30-90%), and canonical vertebrate telomeric repeats (TTAGGG)n counted for 10 kb-long non-overlapping windows (green; 0-300 units), as well as the distribution of tRNA-coding regions (purple) and rDNA loci (red; indicated with an arrowhead).

We compared the Pacific saury genome assembly produced in this study with those of typical teleost fish model species and other beloniform species (Table 2). This comparison highlighted a particularly high proportion of the sequenced genomic regions contained in the chromosome-scale sequences for our Pacific saury assembly (96.8 %). It also showed comparably high coverages (>98 %) of protein-coding landscape of the genome indicated by the number of pre-selected conserved one-to-one orthologs identified, among the species (Table 2). The Pacific saury assembly has fewer gaps (strings of undetermined nucleotide ‘N’ with no less than five bases, in this case) than most of the other chromosome-scale genome assemblies (Table 2). These metrics show a high utility of the Pacific saury genome assembly produced in this study.

The contents of the genomic sequences can be further scrutinized by aligning them with genomic sequences of a closely related sequences. For this purpose, we compared the Pacific saury genome assembly with that of the Japanese medaka^31^ (Fig. 4) which is visualized in a dot plot. This comparison resulted in the regions of cross-species similarity from a chromosome pair residing mostly in a single grid in Figure 4. This suggests high reliability of intra-chromosomal linkages of the Pacific saury genomic contigs that support our chromosome-scale scaffolding as well as high conservation of genomic structure between the two species, with far more intra-chromosomal rearrangements rather than inter-chromosomal ones.

**Figure 4.**
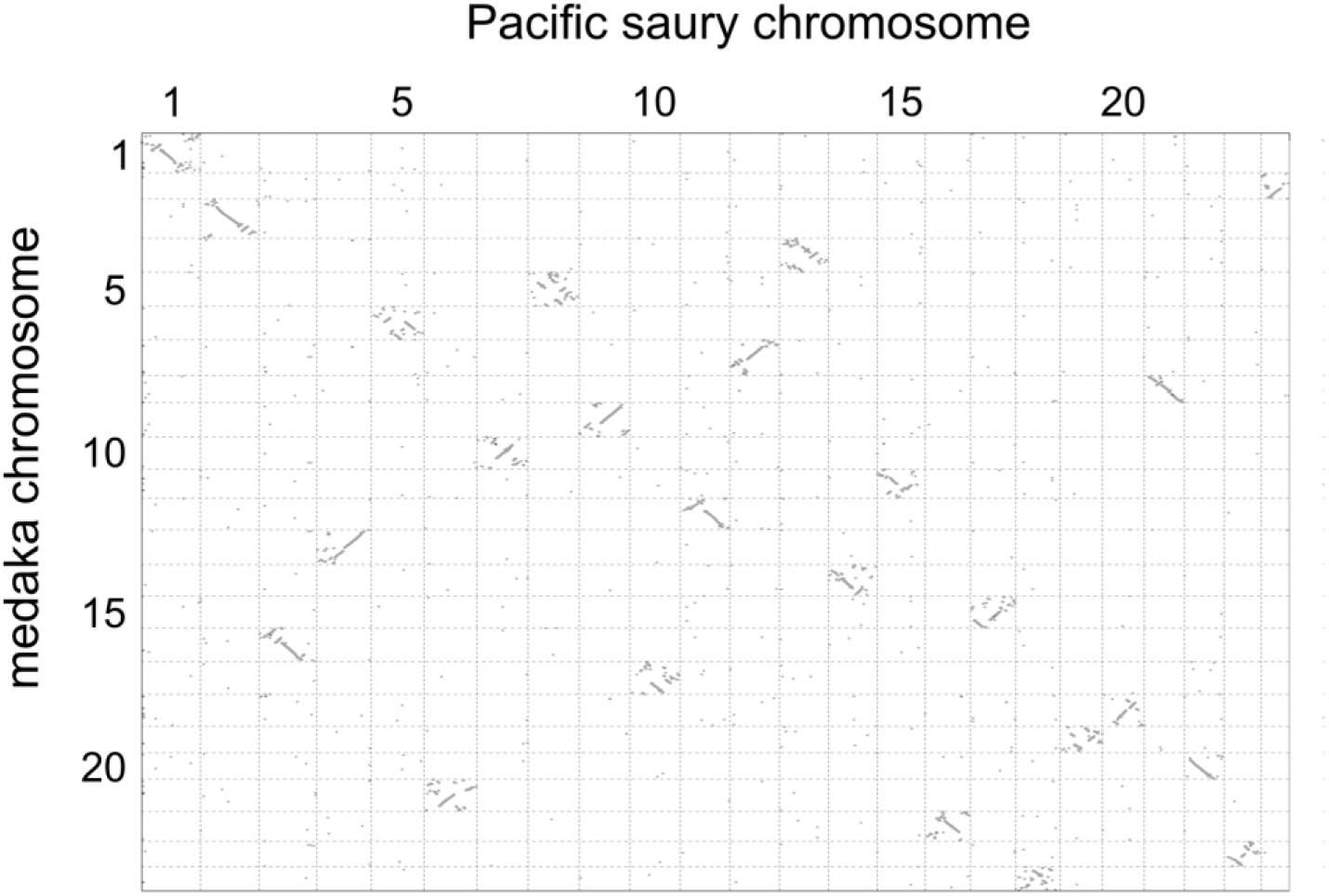
Cross-species comparison of chromosomal sequences. Chromosome-scale sequences of the Pacific saury obtained in this study (total length: 1,111 Mb) and those of the Japanese medaka (*Oryzias latipes*) (total length: 734 Mb) were input as target and query in the D-Genies (https://dgenies.toulouse.inra.fr/) respectively, and the alignment between the two whole genome assemblies was performed with minimap2 (version 2.24). Diagonal lines show regions of continuous similarity.

### 3.4. Non-coding landscape of the genome

Our identification process of repetitive elements, including species-specific (‘*de novo*’) elements, detected repetitive elements in the 51.3% of the whole Pacific saury genome, while those in the medaka genome occupied only 39.5%, based on the same method and criterion. When broken into different repeat classes, the Pacific saury genome showed particularly higher proportion of DNA transposons than that of the medaka (12.9 vs. 3.16 %) (Fig. 5).

**Figure 5.**
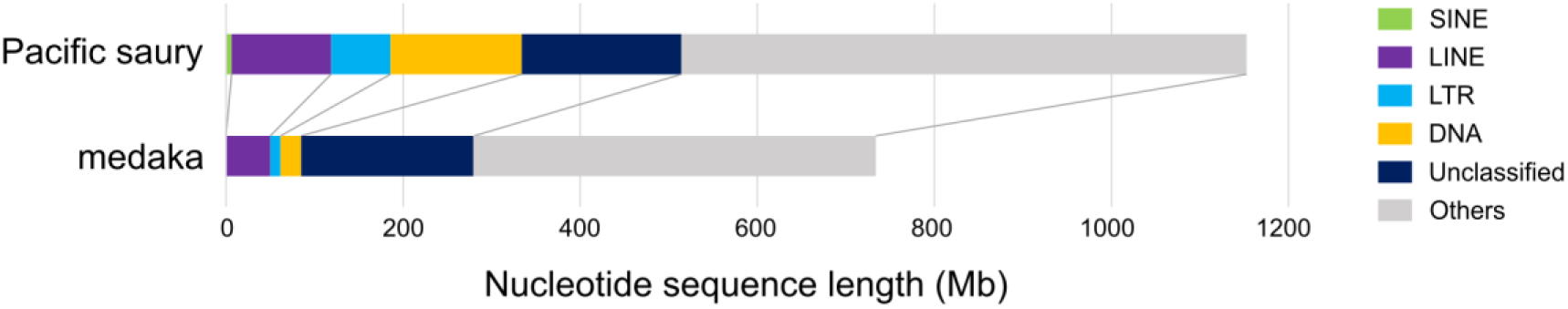
Comparison of repetitive element breakdown into different repeat classes. See Materials and Methods for details. Only chromosome-scale genome sequences were subjected to repeat detection. Distributions of the divergence of detected repetitive sequences are shown in Supplementary Fig. S2.

Chromosome ends consistently exhibited higher GC-content and higher repeat element frequency (Fig. 3B). Among the simple repeats, canonical telomeric repeats (TTAGGG)n were mostly detected closer to, but not always on, the ends of chromosome-scale scaffold sequences (Fig. 3B), which necessitates in-depth investigation to examine whether the chromosome ends are undergoing any lineage-specific structural modification or includes any misjoins in the genome assembly.

We also performed a genome-wide scan of genomic segments containing ribosomal DNA (rDNA) whose intricate genomic structure was one of the major confounding factors in completing the human genome sequencing^32,33^. This analysis was aided by the existing sequence entry OP151180.1 partially covering a Pacific saury rDNA region (6,848 bp) in the NCBI Nucleotide database. Our search detected two units of a 23 kb-long string of 28S, 5.8S, and 18S rDNA genes on chromosome 15 (Fig. 6). We observed more regions with high similarity, but none of those with >90 % sequence identity was included in chromosome-scale sequences (scaffold 1-24). The retrieval of the rDNA loci in the obtained Pacific saury genome assembly shows a higher completeness of the non-coding landscape than the currently available medaka genome assembly (GCF_002234675.1) in which no intact 28S, 5.8S, and 18S rDNA gene cluster is included in its chromosome-scale sequences. Still, the exact number of repetitive units of the rDNA loci remains to be rigidly confirmed with longer sequence reads.

**Figure 6.**
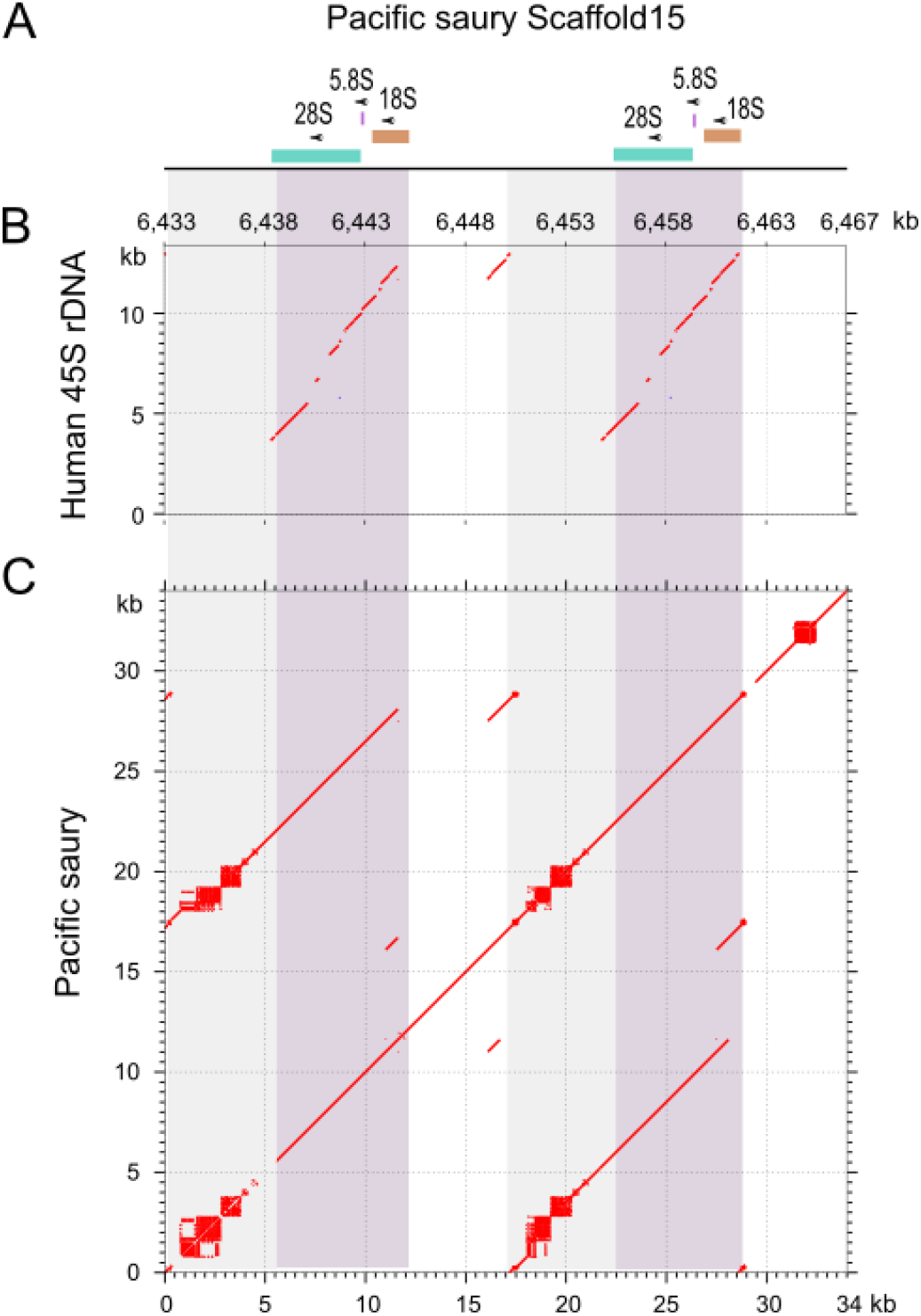
Ribosomal DNA loci in the Pacific saury genome. The 34 kb-long genomic region (position 6,433,000-6,467,000 on Scaffold15) containing two units of the ribosomal DNA cluster (28S-5.8S-18S) is shown. (A) Location of individual rDNA loci. (B) Dot matrix comparing the rDNA-containing Pacific saury genomic region and the human 45S pre-rDNA sequence (NR_145819.1). (C) Dot matrix comparing the rDNA-containing Pacific saury genomic region with itself to indicate internal repeats in it. Light purple background shows an approximately 14 kb-long rDNA-containing regions, and grey background shows the rest of the repetitive unit.

Our assessment of non-coding landscape of the genome encompassed the census of transfer RNAs (tRNAs) whose abundance and distribution in vertebrate genomes are rarely studied (see ^34,35^). Our search, based on both sequence similarity and secondary structure, detected 11,088 candidates for tRNA decoding 20 standard amino acids in the Pacific saury genome, while the medaka genome harbored only 1,894 on the same detection method and criterion. This discrepancy demands in-depth comparative analysis including more species to characterize its biological significance.

### 3.5. Inference of transcribed and translated genomic regions

Animal dissection was performed for a different individual of unknown sex to sample the heart, muscle, gill, liver, gut, and eye ball of an adult as well as a whole larvae body (Supplementary Table S3). The RNAs extracted from these tissues were subjected to library preparation for short-read sequencing. We performed a comprehensive detection of protein-coding regions in the obtained genome assembly with the program pipeline Braker3 using an alignment of the transcriptome short reads from adult and larvae Pacific saury tissues as evidence (398 million read pairs in total) as well as an exploitative protein sequence data set as hints (see Materials and Methods). This resulted in 22,274 predicted protein-coding genes, which are contained in a ready-to-use gene model file that is released under our original website https://treethinkers.nig.ac.jp/saira/. Of these predicted genes, only 24 were predicted outside the 24 chromosome-scale scaffolds.

We compared the protein-coding landscape of the Pacific saury genome with a close relative medaka, with reference to a remarkable difference of the genome size (1.17Gb versus 0.80 Gb^31^) between the two species. On the ground of similar numbers of protein-coding genes (22,274 versus 22,059), the Pacific saury had comparable mean gene length and mean intron length (Table 3). This suggests that the difference in the genome size is attributed to uncharacterized difference outside protein-coding regions.

**Table 3.** Pacific saury genome annotation and its cross-species comparison.

| Feature | Pacific saury | Medaka |
| --- | --- | --- |
| Number of protein-coding genes | 22,266 | 22,059 |
| Mean gene length (nt) | 17,820 | 20,607 |
| Mean intron length (nt) | 1,888 | 1,886 |
| Longest gene (nt) | 488,497 | 1,049,862 |
| Longest cds (nt) | 26,661 | 94,659 |
| Longest exon (nt) | 13,178 | 19,832 |
| Longest intron (nt) | 459,515 | 1,035,396 |
| Number of predicted tRNA<sup>1</sup> | 11,088 | 1,894 |
| Repetitiveness (%)<sup>2</sup> | 51.3 | 39.5 |

The statistics for medaka was computed for the gene model in the .gff file registered for the genome assembly GCF\_002234675.1<sup>31</sup>

### 3.6. Census of aquaporin (AQP) gene family members

We scanned the obtained genome assembly to identify the coding genomic regions of the genes encoding aquaporin (AQP) proteins that function mainly as water channels^36^. The coverage of AQP genes, often intervened by long introns, is expected to allow an additional metric for the evaluating the continuity of genomic sequences. Our search identified 15 Pacific saury AQP genes, and they were categorized into eleven major subgroups that presumably existed in the common jawed vertebrate ancestor (Fig. 7) by our molecular phylogeny inference (Supplementary Fig. S3). Because of the teleost-specific whole genome duplication^37^, two paralogs were identified for AQP0, -9, -10, and -11, as previously shown for some teleost fishes^38^. In Pacific saury genome, we could not detect the multiplicity of the AQP1, -3, and -8 which was previously observed in the zebrafish but not in the medaka, suggesting that their absences in the medaka is not due to false negative detection. The retrieval of all the AQP genes possessed by the medaka that belongs to the same taxon Beloniformes ascertains the high completeness of the Pacific saury genome assembly obtained in this study.

**Figure 7.**
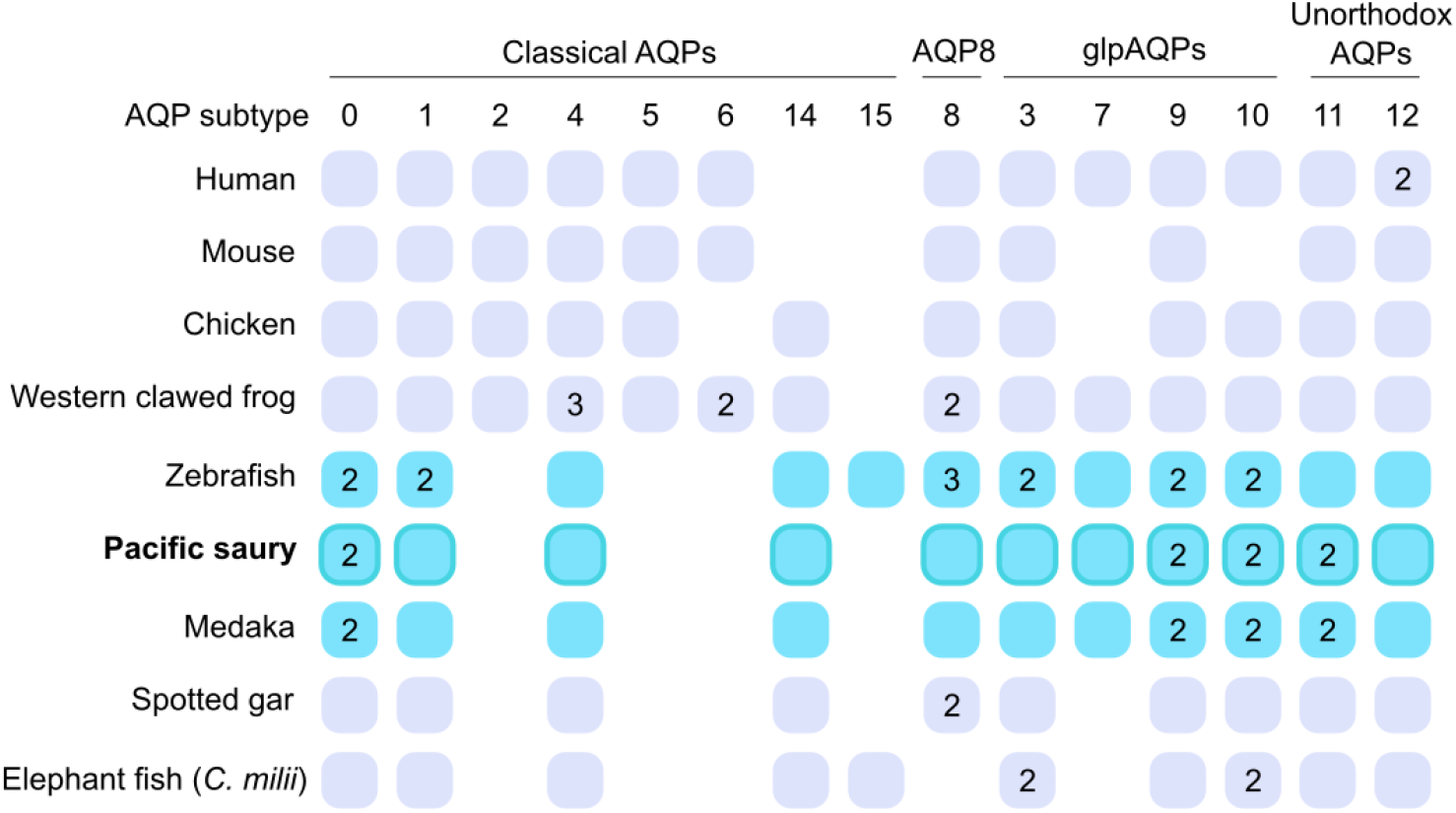
Aquaporin (AQP) gene identification. Orthology-based schematic categorization of Pacific saury genes. Different AQP subtype genes are positioned based on the orthology suggested by the molecular phylogenetic tree in Supplementary Fig. S3. The numbers in the boxes indicate the multiplicity of the orthologs generated by lineage-specific gene duplications, while no box shows the absence of ortholog in the currently available genome assemblies. Telest fish gene repertoires are highlighted in cyan. At the top is the classification into different subfamilies including glpAQPs (aquaglyceroporins) based on existing literature^36,39^.

## Acknowledgements

We thank Mayuki Ando and Hiroaki Chiba for their helpful advice and assistance during the histological experiments, Miyuki Mekuchi, Yoji Nakamura, and Motoshige Yasuike at FRA for insightful discussion about fishery science of the Pacific saury, Yoshinobu Uno for discussion on karyotyping reports, Osamu Nishimura for quality control of short read sequencing data, Yoshinori Hasegawa for conducting HiFi sequence data production, and Rintaro Ishii, Toshiaki Mori, Hikari Yoshizato for their maintaining fish colonies.

## Funding

This research was supported by “Strategic Research Projects” grant from ROIS (Research Organization of Information and Systems).

## Conflict of interest

None declared

## Data availability

Raw sequence reads and the genome assembly have been deposited in NCBI under the BioProject ID PRJNA1027303. The predicted coding sequences (in both nucleotides and amino acids) and the gene models (in the .gff3 file format), as well as the genome assembly, are also available at https://figshare.com/projects/Pacific-saury-genome/178218.

## Supplementary Information

**Supplementary Figure S1.**
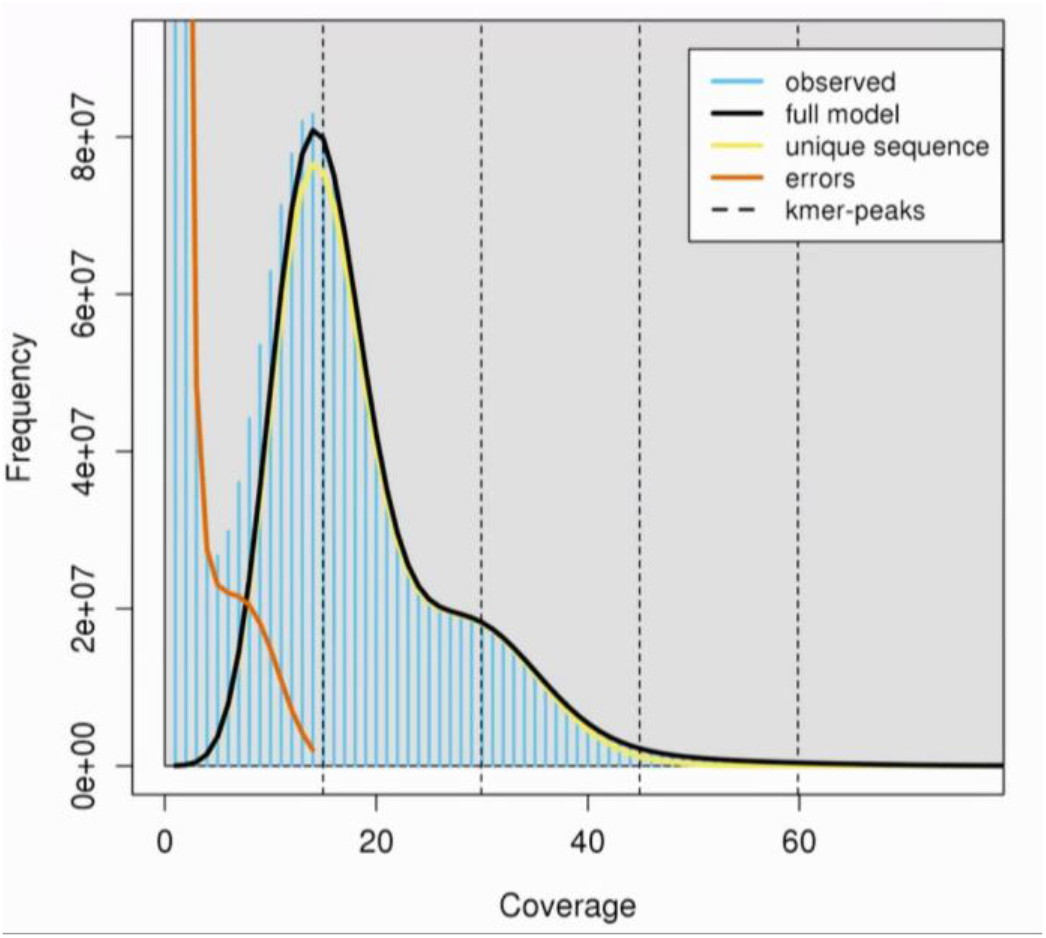
Distribution of k-mers based on HiFi reads. K-mer frequency was computed with the jellyfish v2.3.0, and the resultant .histo file is input in the GenomeScope^40^.

**Supplementary Figure S2.**
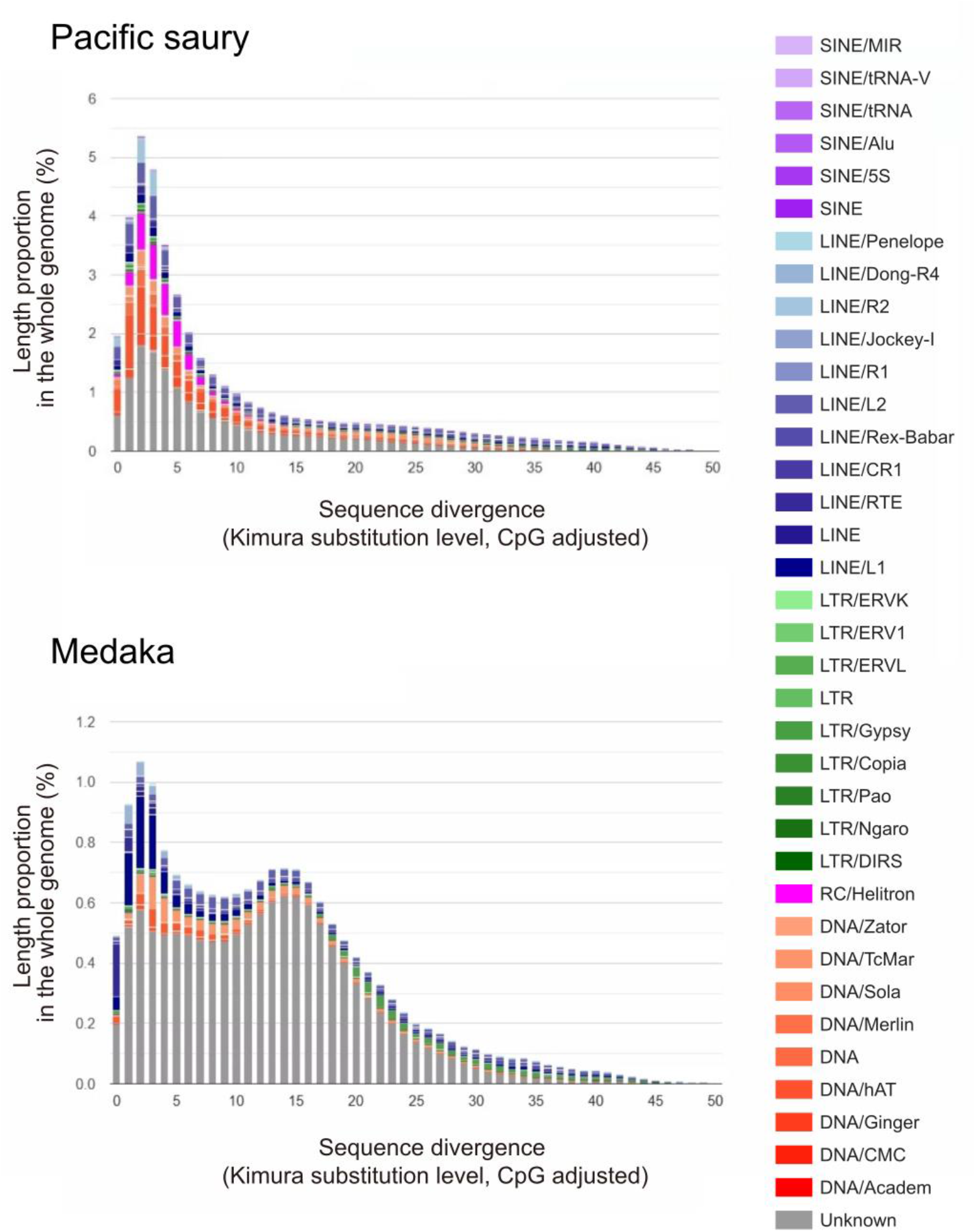
Repetitive element compositions and their divergence profiles. Distribution of sequence divergence of the identified repetitive elements. The divergence of repetitive elements in individual repeat subclasses was computed using the Perl scripts, calcDivergenceFromAlign.pl and createRepeatLandscape.pl, that are accompanying the program RepeatMasker. See Materials and Methods for details of repeat detection. Note that the height is not equally scaled between the species.

**Supplementary Figure S3.**
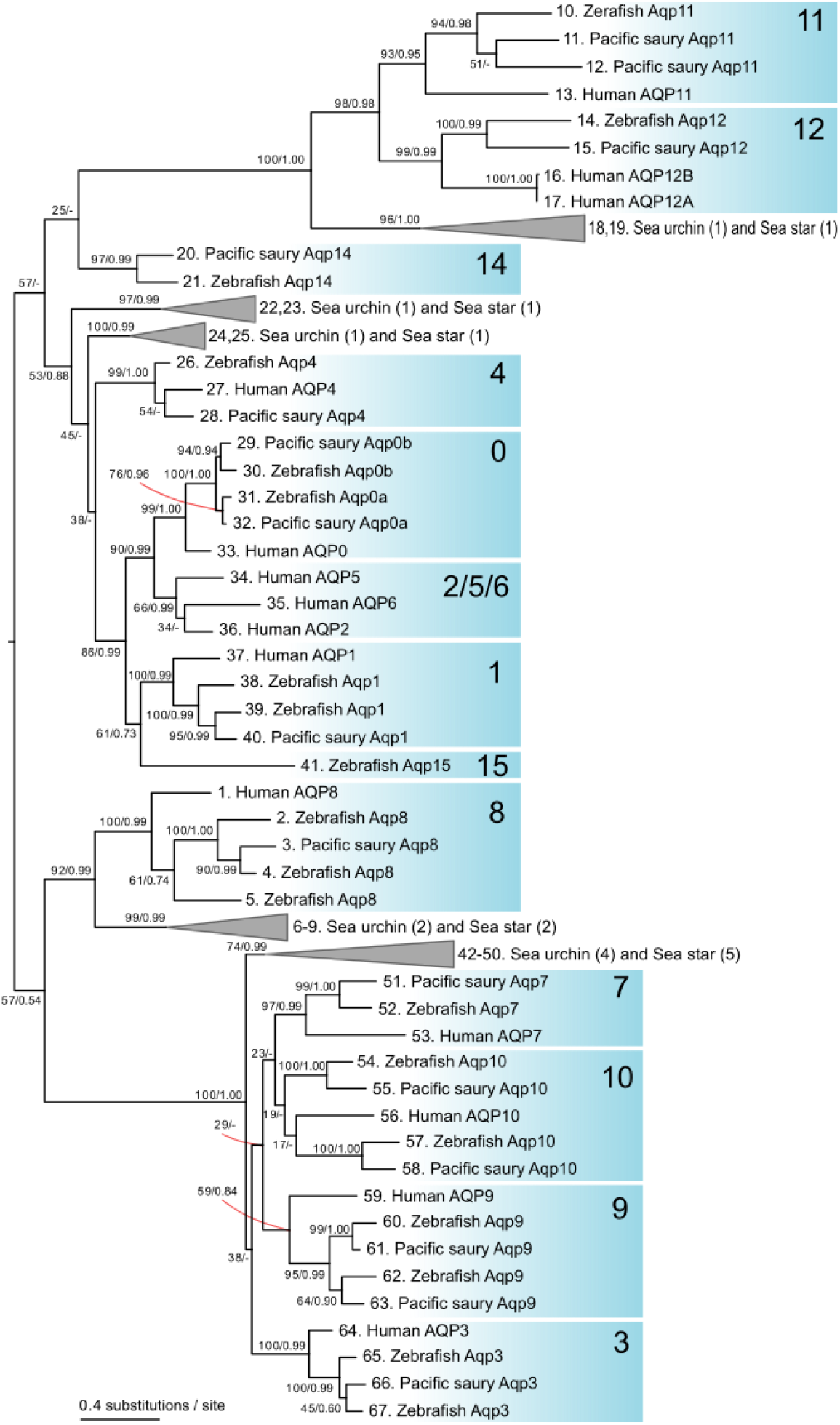
Aquaporin gene phylogeny. Phylogenetic tree confirming the orthology of the Pacific saury genes. The maximum-likelihood tree was inferred with 232 residues in the amino acid sequence alignment (see Materials and Methods). The support values at notes are bootstrap probabilities in the ML tree and posterior probabilities in the Bayesian inference (see Materials and Methods). The numbers in the brackets shown the multiplicity of the sequences used for the species indicated.

**Supplementary Table S1.**
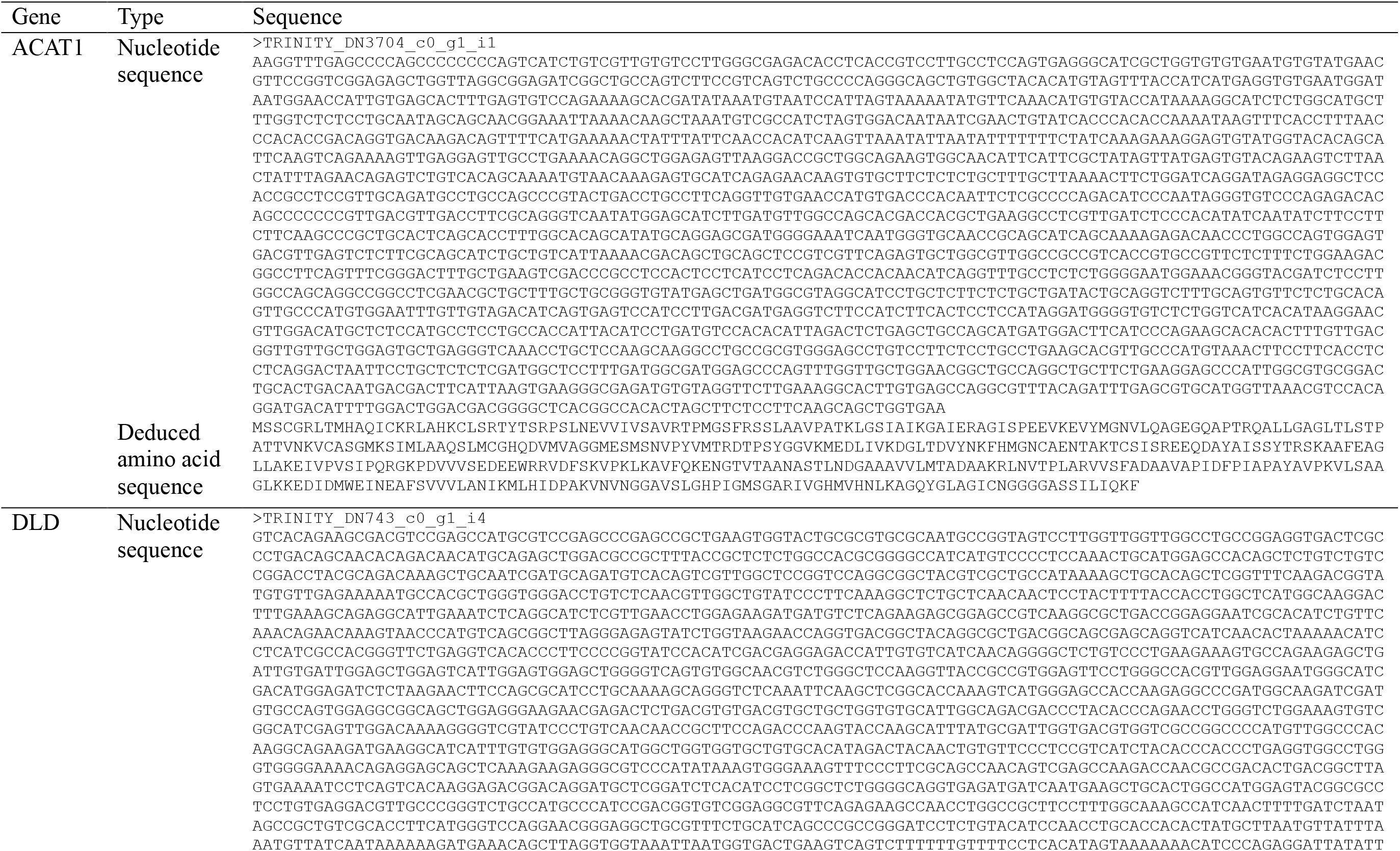
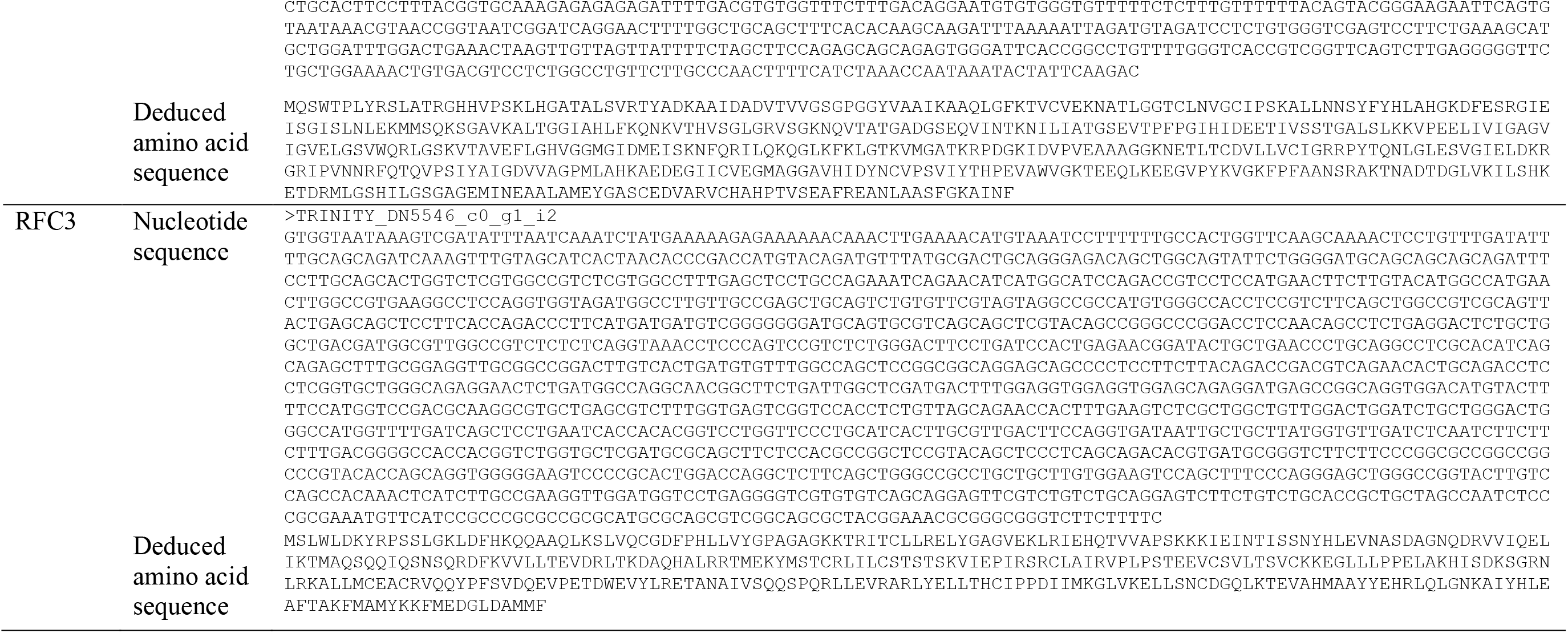
Nucleotide and deduced amino acid sequences of the transcripts used for the analysis.

**Supplementary Table S2.**
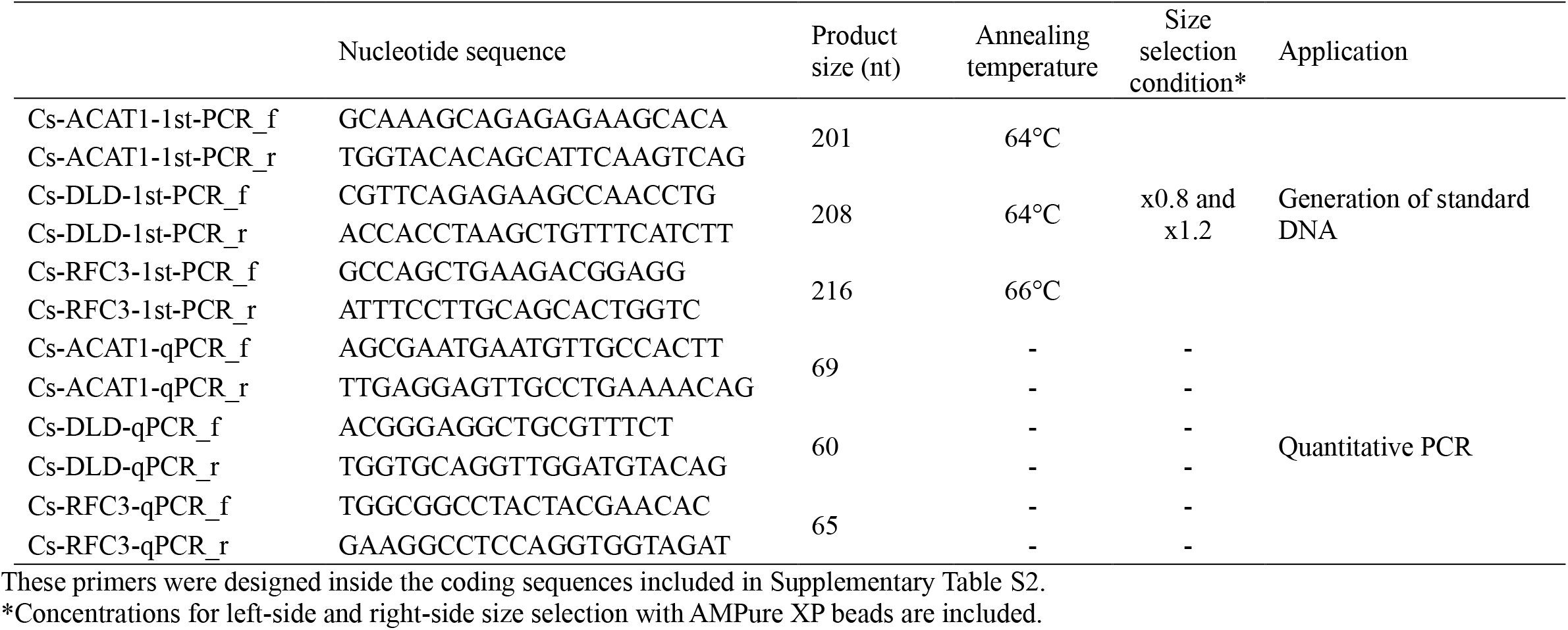
Oligonucleotide primer sequences and DNA preparation conditions.

**Supplementary Table S3.** RNA-seq library preparation and sequencing.

| Library ID | Tissue | RIN | Number of PCR cycles | Number of raw reads obtained (million read pairs) |
| --- | --- | --- | --- | --- |
| P596_01_1 | adult eyeball | 9.4 | 11 | 114.4 |
| P596_02_1 | adult heart | 8.9 | 11 | 121.8 |
| P596_03_1 | adult gill | 8.4 | 11 | 119.5 |
| P596_04_1 | larvae whole | 9.8 | 11 | 108.5 |
| P596_05_1 | adult muscle | 9.1 | 11 | 101.0 |
| P596_06_1 | adult liver | 7.5 | 12 | 109.4 |
| P596_07_1 | adult gut | N/A | 12 | 121.0 |

**Supplementary Table S4.** Aquaporin (AQP) gene entries used of molecular phylogeny inference.

| # in tree | Orthogroup* | species | Accession ID |
| --- | --- | --- | --- |
| 1 | AQP8 | *Homo sapiens* | NP_001160.2 |
| 2 | AQP8 | *Danio rerio* | ACV60540.2 |
| 3 | AQP8 | *Cololabis saira* | This study |
| 4 | AQP8 | *Danio rerio* | NP_001073651.1 |
| 5 | AQP8 | *Danio rerio* | NP_001004661.1 |
| 6 | AQP8 | *Strongylocentrotus purpuratus* | XP_030854916.1 |
| 7 | AQP8 | *Patiria miniata* | XP_038075258.1 |
| 8 | AQP8 | *Patiria miniata* | XP_038069095.1 |
| 9 | AQP8 | *Strongylocentrotus purpuratus* | XP_030840396.1 |
| 10 | AQP11 | *Danio rerio* | AAH95775.1 |
| 11 | AQP11 | *Cololabis saira* | This study |
| 12 | AQP11 | *Cololabis saira* | This study |
| 13 | AQP11 | *Homo sapiens* | NP_766627.1 |
| 14 | AQP12 | *Danio rerio* | AAI21753.1_ |
| 15 | AQP12 | *Cololabis saira* | This study |
| 16 | AQP12 | *Homo sapiens* | NP_001095937.1 |
| 17 | AQP12 | *Homo sapiens* | NP_945349.1 |
| 18 | AQP11 | *Patiria miniata* | XP_038044509.1 |
| 19 | AQP11 | *Strongylocentrotus purpuratus* | XP_003725070.1 |
| 20 | AQP14 | *Cololabis saira* | This study |
| 21 | AQP14 | *Danio rerio* | XP_005174182.1 |
| 22 | cAQP | *Patiria miniata* | XP_038051349.1 |
| 23 | cAQP | *Strongylocentrotus purpuratus* | XP_001190612.2 |
| 24 | cAQP | *Patiria miniata* | XP_038078625.1 |
| 25 | cAQP | *Strongylocentrotus purpuratus* | XP_001185961.1 |
| 26 | AQP4 | *Danio rerio* | XP_009292895.1 |
| 27 | AQP4 | *Homo sapiens* | XP_011524244.1 |
| 28 | AQP4 | *Cololabis saira* | This study |
| 29 | AQP0 | *Cololabis saira* | This study |
| 30 | AQP0 | *Danio rerio* | NP_001018356.1 |
| 31 | AQP0 | *Danio rerio* | NP_001003534.1 |
| 32 | AQP0 | *Cololabis saira* | This study |
| 33 | AQP0 | *Homo sapiens* | NP_036196.1 |
| 34 | AQP5 | *Homo sapiens* | NP_001642.1 |
| 35 | AQP6 | *Homo sapiens* | NP_001643.2 |
| 36 | AQP2 | *Homo sapiens* | NP_000477.1 |
| 37 | AQP1 | *Homo sapiens* | NP_932766.1 |
| 38 | AQP1 | *Danio rerio* | XP_021336241.1 |
| 39 | AQP1 | *Danio rerio* | NP_996942.1 |
| 40 | AQP1 | *Cololabis saira* | This study |
| 41 | AQP15 | *Danio rerio* | XP_021327889.1 |
| 42 | glpAQP | *Patiria miniata* | XP_038071748.1 |
| 43 | glpAQP | *Patiria miniata* | XP_038073806.1 |
| 44 | glpAQP | *Patiria miniata* | XP_038073167.1 |
| 45 | glpAQP | *Strongylocentrotus purpuratus* | XP_792142.4 |
| 46 | glpAQP | *Strongylocentrotus purpuratus* | XP_030833261.1 |
| 47 | glpAQP | *Strongylocentrotus purpuratus* | XP_789770.3 |
| 48 | glpAQP | *Patiria miniata* | XP_038073505.1 |
| 49 | glpAQP | *Patiria miniata* | XP_038071359.1 |
| 50 | glpAQP | *Strongylocentrotus purpuratus* | XP_030832856.1 |
| 51 | AQP7 | *Cololabis saira* | This study |
| 52 | AQP7 | *Danio rerio* | NP_956204.2 |
| 53 | AQP7 | *Homo sapiens* | NP_001161.1 |
| 54 | AQP10 | *Danio rerio* | ACB10577.1 |
| 55 | AQP10 | *Cololabis saira* | This study |
| 56 | AQP10 | *Homo sapiens* | NP_536354.2 |
| 57 | AQP10 | *Danio rerio* | NP_001002349.1 |
| 58 | AQP10 | *Cololabis saira* | This study |
| 59 | AQP9 | *Homo sapiens* | NP_066190.2 |
| 60 | AQP9 | *Danio rerio* | NP_001028268.1 |
| 61 | AQP9 | *Cololabis saira* | This study |
| 62 | AQP9 | *Danio rerio* | NP_001171215.1 |
| 63 | AQP9 | *Cololabis saira* | This study |
| 64 | AQP3 | *Homo sapiens* | NP_004916.1 |
| 65 | AQP3 | *Danio rerio* | NP_998633.1 |
| 66 | AQP3 | *Cololabis saira* | This study |
| 67 | AQP3 | *Danio rerio* | NP_001159593.1 |

\* Note that the gene product name in NCBI are often different from the gene name in this column which was inferred with molecular phylogeny inference.

## Notes

https://figshare.com/projects/Pacific-saury-genome/178218

### Competing Interest Statement

The authors have declared no competing interest.

